# Age-Group Differences in Skeletal Age, Anabolic Hormones, and Physical Fitness in Elite Youth Football Players from Under-13 to Under-16

**DOI:** 10.64898/2026.06.11.731548

**Authors:** Ebrahim Eskandarifard, Mehdi Kargarfard, Rafael Oliveira, Seyed Abbas Afsanepurak, Hadi Nobari, António José Figueiredo

## Abstract

**Objectives:** Football players between the ages of 13 and 16 were the subjects of this study to (i) compare skeletal age, growth hormone (GH), insulin-like growth factor-1 (IGF-1), VO_2_max, countermovement jump (CMJ) performance, and playing time across four consecutive age groups (under-13 to under-16), and (ii) investigate correlations between these metrics within each age group.

**Design:** cross-sectional research study.

**Methods:** The U13, U14, U15, and U16 groups included 122 elite male youth football players (14.13 ± 1.14 years; 169.39 ± 9.82 cm; 55.56 ± 10.10 kg). The Fels method was used to determine skeletal age. Anthropometric measures included height, weight, and sitting height. The CMJ and Yo-Yo (VO_2_max) tests were used to assess physical fitness. GH and IGF-1 were measured in blood samples. Pearson correlation and one-way ANOVA were used.

**Results:** Anthropometric assessment, skeletal age, VO_2_max, CMJ, IGF-1, football training, and minutes playing all showed significant group differences, with the U16 group demonstrating the highest values and the U13 group the lowest (p < 0.001). There were no significant differences in GH (p = 0.172). Although there were no consistent relationships between GH or IGF-1 and height or weight, these variables did correlate with other factors; however, these associations varied by group.

**Conclusions:** Across the under-13 to under-16 span, skeletal age, physical fitness, and IGF-1 increased progressively between age groups, whereas GH did not differ. The impact of biological maturation on performance was demonstrated by the fact that older, more experienced players performed better than younger ones. The absence of a between-group difference in GH, despite progressive changes in the other measures, points to maturity-driven adaptations rather than a uniform hormonal response.

## Introduction

Physical activity plays an important role in the growth and development of children and adolescents [1]. In sports such as football, different age categories are related to different physical and physiological responses [2]. For that reason, the assessment of physical and physiological variables is extremely important for understanding those differences and establishing the effectiveness of individual blocks of training or the overall success of the annual training plan [3].

Some measures that are usually quantified in football are related to the anaerobic system, such as jumping, sprinting, or scoring a goal [4]. In addition, football is also related to aerobic fitness, and for that reason, another common variable quantified is the maximum rate of oxygen consumption (VO_2_max), which helps to identify the level of physical and physiological fitness [5-8].

Furthermore, intense sport training and competition are often linked together [9]. In addition to these factors, normal growth in children and adolescents is largely regulated by growth hormone (GH) and insulin-like growth factor 1 (IGF-1) [10]. Exercise plays an important role in regulating the secretion of these two hormones. Research has shown different results in terms of the GH response to exercise [11, 12]. Athletes who overtrain suppress GH production, and their tolerance for exercise decreases, and, eventually, their performance decreases. Thus, the GH response to exercise is complex and appears to be affected by many variables (e.g., type of exercise, intensity, duration, age, etc.) [12]. The effect of exercise on IGF-1 responses remains unclear. Some researchers have reported that endurance training increases the levels of this hormone [13-15]. However, other evidence suggests that IGF-1 levels may decrease under specific exercise conditions, such as acute high-intensity efforts, although findings remain inconsistent across studies [13-15]. On the other hand, professional sports activities seem to have a significant effect on the secretion of various hormones, including GH and IGF-1. For example, Barbara et al. conducted a study in 2020 that was designed to investigate whether exercise-induced increases in testosterone levels affect the growth hormone/insulin-like growth factor (GH-IGF-1) axis and body composition, particularly skeletal muscle mass. They reported that exercise causes significant changes in the cortisol/free testosterone ratio, which can affect the secretion of GH and IGF-1 from the liver and ultimately into lean body mass [16].

More research is needed on young elite players to better identify growth and hormonal changes. Some cross-sectional studies have shown that football training promotes positive hormonal adaptation [17, 18]. Nevertheless, the effects of long-term and intense football training on GH/IGF-I hormonal compliance are rare. Only one study has examined the effects of intense football training during a competitive season on growth hormones [17]. Hammami et al. (2018) studied the effects of two seasons of football training on the growth, development, and concentration of somatotype hormones in elite youth football players. The results of their study revealed that intensive training does not harm the growth or development of young football players for 2 years [12]. It seems that the study of growth factors in young football players is highly important. Therefore, the positive and negative features of training at a young age should be well defined.

Football is the most popular sport in the world, especially among children and teenagers. Optimizing the physical potential of young football players is one of the main goals of football academies [19, 20]. In fact, elite football players need to be prepared for strenuous training. The most important dimensions for measuring performance in football are physical fitness and technical and tactical performance [12, 20]. The physical fitness of football players is usually measured in terms of endurance, speed, strength, and power [19]. It is relatively easy to assess the physical fitness of young players, but it is more difficult to distinguish between adapting to football training and developing mediators [18].

Another major assessment in young players is maturation and skeletal age. Skeletal age, which is predicted by hand radiography, is considered to be the most appropriate method for estimating biological maturity [21]. Maturation, which refers to the progress of the mature state, can vary in timing and between different body systems [22]. Different young players could be biologically ahead of their chronological age (early maturing individual), equal to their chronological age, or they could stay behind their chronological age (late-maturing individual) [23]. Therefore, physiological monitoring at such young ages is important and recommended. With the information of biological maturity, training periodization and prescription could be better adjusted in terms of physiological and physical knowledge [24]. Although maturation, physical fitness, and hormonal indices have been examined previously in elite youth footballers, prior work has typically done so within a single age category (under-15 or under-16) and has focused mainly on explaining individual variation in playing time [25-28]. How these maturational, hormonal, and fitness characteristics differ between consecutive age groups across the under-13 to under-16 span, within one academy and using a common assessment protocol, remains unclear. Characterizing these between-group differences situates each age category along a developmental continuum rather than in isolation, and may help academies calibrate expectations and training prescription as players progress from under-13 to under-16.

In Iran, the skeletal age of young elite players is not well defined. Given that there are only a few well-known football academies in Iran where elite players are presented, one of these academies was used to compare skeletal age, anabolic hormones, and physical fitness between consecutive age groups of elite football players from under-13 to under-16. Therefore, this study had two purposes: (i) to investigate differences between age (under 13 to under 16) groups in skeletal age, VO_2_max, countermovement jump (CMJ) performance, and GH and IGF1 levels and playing time; and (ii) to elucidate the relationships among the previous measures within each age group. We predicted that older players (U16), because of their advanced biological maturation and greater training experience, would have higher values in anthropometric measures (height, weight, sitting height, and leg length), skeletal age, VO_2_max, CMJ performance, and IGF-1 levels than younger players (U13) would have on the basis of prior research [29-31]. However, given that previous research indicates that GH responses to exercise may be less influenced by age in young athletes [12, 17], we did not anticipate any substantial changes in GH levels across groups. Furthermore, owing to the complicated control of the GH-IGF-1 axis during adolescence, we predicted that skeletal age, playing time, and physical fitness measurements (VO_2_max and CMJ) would be positively linked with anthropometric traits and IGF-1 levels but not necessarily with GH [14, 32].

## Methodology

### Participants

The participants in this study included 122 elite youth football players (mean ± standard deviation (SD)); age: 14.13 ± 1.14 years; height: 169.39 ± 9.82 cm; weight: 55.56 ± 10.10 kg; skeletal age: 14.94 ± 1.42 years; VO_2_max: 47.22 ± 3.76 ml·kg^−1^·min^−1^) (Table 1). All players were divided according to their age category: U13 (n=30), U14 (n=31), U15 (n=30), and U16 (n=31). They all played at the highest level of their age group. Only U16 played in the national league, while the other age groups played in the provincial league. The football training experience for each group was as follows: U13, 5.25 ± 1.60 years; U14, 5.62 ± 1.87 years; U15, 6.69 ± 1.80 years; and U16, 6.65 ± 1.55 years. This study was authorized by the ethics committee of the University of Isfahan by the code of IR.UI.REC.1399.001 and was developed according to the Helsinki Declaration. All the participants and their parents were informed about the risks and benefits of this study. They had the right to quit in each part of that they wanted, and as all participants were minors, written informed consent was obtained from their parents or legal guardians and this form signed as a testimonial to this study. The inclusion criteria were as follows: 1) had a history of attending the club academy in the lower ranks for at least 4 years; 2) cooperated for regular participation in all stages of the study; 3) did not use any growth stimuli under the supervision of a specialist doctor or the relevant club; and 4) were not allowed to perform extra exercise. The exclusion criteria for this study were as follows: 1) did not participate in 80% of the exercises during the whole season and 2) did not participate in one of the medical tests or physical evaluations of the study.

**Table 1:** ANOVA Analysis of Players in Different Groups.

| Variables | U13 (n=30) | U14 (n=31) | U15 (n=30) | U16 (n=31) | F | P |
| --- | --- | --- | --- | --- | --- | --- |
|  | Mean ± SD | Mean ± SD | Mean ± SD | Mean ± SD |  |  |
| Height (cm) | 158.52 ± 8.64 | 169.84 ± 6.98 | 172.41 ± 7.59 | 176.53 ± 5.66 | 33.92 | <0.001 |
| Weight (kg) | 46.37 ± 6.72 | 55.31 ± 7.98 | 59.48 ± 10.32 | 60.92 ± 8.49 | 18.02 | <0.001 |
| Sitting Height (cm) | 79.73 ± 4.53 | 85.80 ± 3.66 | 89.98 ± 3.34 | 93.01 ± 2.40 | 79.10 | <0.001 |
| Leg Length (cm) | 78.78 ± 5.16 | 84.04 ± 4.35 | 82.42 ± 4.98 | 83.52 ± 4.26 | 7.69 | <0.001 |
| Skeletal Age (yrs) | 13.35 ± 0.85 | 14.71 ± 0.62 | 15.01 ± 0.99 | 16.64 ± 0.79 | 82.00 | <0.001 |
| Football Training (mo) | 63.03 ± 19.19 | 67.45 ± 22.39 | 80.23 ± 21.65 | 79.81 ± 18.56 | 5.48 | <0.001 |
| VO<sub>2</sub>max (ml·kg<sup>-1</sup>·min<sup>-1</sup>) | 43.27 ± 1.71 | 46.99 ± 3.08 | 48.71 ± 2.71 | 49.81 ± 3.58 | 30.20 | <0.001 |
| CMJ (cm) | 29.67 ± 3.48 | 32.29 ± 4.44 | 39.47 ± 5.69 | 40.87 ± 3.87 | 45.51 | <0.001 |
| GH (ng/dl) | 0.73 ± 1.29 | 1.36 ± 2.48 | 1.17 ± 1.30 | 1.78 ± 2.01 | 1.69 | 0.172 |
| IGF1 (ng/dl) | 391.07 ± 170.69 | 411.71 ± 139.09 | 457.90 ± 111.86 | 356.32 ± 100.71 | 3.09 | <0.030 |
| Minutes of Playing (min) | 111.16 ± 42.72 | 435.16 ± 248.72 | 1360.50 ± 271.98 | 1383.74 ± 248.98 | 259.78 | <0.001 |

Abbreviations: $VO_{2max}$ : maximal oxygen consumption; CMJ: countermovement jump; GH: growth hormone; IGF1: insulin-like growth factor 1; Yrs: years; Mo: months.

### The Experimental Approach to the Problem

This research was conducted as a quasiexperimental and cohort study that was performed on a cross-sectional basis, yielding practical results. Researchers checked players throughout the whole season, and assessments were performed upon completion of the competitive season. Data were collected over one full competitive season, from 1 July 2019 to 30 April 2020. All assessments were performed at the end of the season during the postseason phase (April 2020). At the end of the training season and after three days of recovery from the last training, players underwent anthropometric assessments, hormonal tests, and imaging of left-hand radiography on day 1; the CMJ was evaluated on day 2; and the Bangsbo curved anaerobic test and Yo-Yo Intermittent Recovery Test Level 1 (YIRT1) were tested on the last day. All tests were performed in the morning under the same conditions (21–23°C and 50% humidity) [33].

### Anthropometric Measurements

For anthropometric measurements, three variables (weight, standing height, and sitting height) were assessed, and all measurements were carried out in the morning, in a fasted state (minimum of 8 h) with an empty bladder and without engaging in exercise, drinking alcohol, or consuming any caffeine type in the 24 h prior to the measurements. All the measurements were conducted by the same researcher to minimize possible errors, and the order of the measurements was the same for all the participants. A SECA audiometer (2013, Hamburg, Germany) was used to measure standing height and sitting height. The participants stood as close as possible to the stadiometer with their bare feet, and their head, back, and shoulder were attached to the stadiometer with their feet positioned beside each other to assess standing height. For sitting height, they sat on a 50 cm box with their buttocks as close as possible to the stadiometer, and their posture was upright. The difference between the stadiometer number and the box height is the sitting height; also, to estimate the maturity offset, we need the leg length, which is calculated as the difference between the standing height and the sitting height. The last assessment in this part was the weight that was evaluated by SECA, model 813, England, with an accuracy of ± 0.1 kg [34].

### Skeletal Age

Skeletal age is the most acceptable method for assessing maturity level. Two-dimensional radiographs of the hand wrist were used to verify skeletal age. For this purpose, we used the EOS Imaging system EOS 2D/3D, EOS Imaging, Paris, France, to specify the skeletal age. The EOS imaging system is a new device that can provide a lower radiation dose (50–85%) than digital radiography can provide [35-37], as well as better image quality [38, 39]. There are several techniques for assessing skeletal age, but the most reliable is the Fels method, which we used in this study by our expert [40]. In the Fels method assessment, we used several criteria related to the level of maturity and ratio of hand-wrist bones. Additionally, chronological age, which is the difference between the date of assessment and the birth date, was used. After the ratios and maturity levels of the bones were identified, the data and chronological ages were input into Felshw 1.0 software to evaluate the skeletal age. To identify the maturity status of the player, we had to subtract the skeletal age from the chronological age. If this number was greater than +1, the player was identified as early.

### Blood analysis

After at least 12 hours of fasting and 72 hours after the last training, the players returned to the Al-Zahra Hospital laboratory for the collection of 10 ml of blood. Blood was drawn from the antecubital fossa. This blood was drawn at 8 am, and the samples were quickly centrifuged. To quantify GH and IGF-1 levels, the laboratory used serum, which was expelled from the blood. In this part, the chemiluminescence technique (ICMA) and the IMMULITE framework (2000xpi Systems, organization SIEMENS Germany) were used. The sensitivity of the kit used for GH (REF: L2KGRH2; lot 171) was 0.01 ng/ml, and that used for IGF-1 (REF: L2KGF2 and lot: 571) was 13.3 ng/ml.

### Countermovement Jump Test

The CMJ test was used to investigate explosive lower-body power [41]. The players warmed for approximately 15 min, starting by running slowly, followed by a 5*10 meter run with maximum speed, horizontal and vertical hopping, and countermovement jump drills. Players, before performing the CMJ test, carried out 2 trials for familiarizing. The first test started when the examiner said “jumped,” and the player stood on the mat with 90-degree flexion in the knee; then, they jumped vertically with maximum energy. The second test was performed after 5 min of recovery. Finally, the best performance was recorded in centimeters [42].

### Yo‒Yo Intermittent Recovery Level 1 Test

The aerobic test was the last test performed by researchers, and YIRT1 was used to evaluate VO_2_max. A conditioning coach performed a standard warm-up, and the procedure was similar to the 7 repeated sprint tests (7RST test). In YIRT1, the participants ran 40 meters back and forth, after which they recovered 10 meters back and forth. The test is conducted at 10 km/h and increases by 0.5 km/h for the next step. The end of this test occurred when each of the participants experienced two failures because they were in line at the same time as the beep sound. The VO_2_max was evaluated via the following equation: VO_2_max (ml·kg^−1^·min^−1^) = YRIR1 distance (m) × 0.0084 + 36.4 [42].

### Sport-Specific Training Age

In the scientific literature, the term “sport-specific training” is used, and since this research focuses on football, we look specifically at the length of time each player has been involved in football. This duration, measured in months, represents the total time a player has been actively playing football since the beginning of their career. To gather this information, we reviewed each player’s history via the club’s archive and database. For greater accuracy, we then cross-checked these results with the players themselves, recording the final data as “football training” in months [27, 43]. On the other hand, the training duration for each player, representing the total months of football-specific training since the beginning of their career, was collected from the academy’s records and verified with the players. The values are reported as averages for each group.

### Statistical Analyses

Statistical analyses were performed via SPSS version 22 (IBM Corp, Armonk, NY, USA). First, the groups were characterized through descriptive statistics by means and SDs (standard deviations). Second, Shapiro‒Wilk and Levene tests were conducted to determine normality and homoscedasticity, respectively. Once variables obtained a normal distribution (Shapiro‒Wilk > 0.05), ANOVA and the Bonferroni post hoc correction were used to compare all groups at a significance level of P < 0.05. Third, Pearson correlation analysis was performed between minutes of playing, physical fitness test results, IGF1, GH, and skeletal age. The correlation magnitudes were classified as follows: r < 0.3 (weak), 0.3–0.5 (moderate), 0.5–0.7 (strong), and >0.7 (very strong) [44]. Finally, Cohen’s d effect size (ES) statistic was calculated to analyze the magnitude of effects of the ANOVA comparisons through the difference in means divided by the standard deviation of the data. According to Hopkins [45], the following threshold scale was applied: <0.2 = trivial, 0.2–0.6 = small, 0.6– 1.2 = moderate, 1.2–2.0 = large, and >2.0 = very large effect.

## Results

The descriptive characteristics of the players are presented in Table 1, where values are reported as the means ± SDs for each group (U13, U14, U15, U16) rather than as a whole-group average. The total season consisted of 824.06 ± 603.53 minutes of playing across all groups. The mean chronological age for U13 was 12.61 ± 0.29 years, that for U14 was 13.65 ± 0.24 years, that for U15 was 14.62 ± 0.23 years, and that for U16 was 15.61 ± 0.25 years.

The results of the ANOVA shown in Table 1 revealed that there were significant differences between groups in terms of anthropometric characteristics (height, weight, sitting height, and leg length), skeletal age, VO_2_max, CMJ, IGF-1, football training, and minutes of playing, but there were no significant differences in GH levels. To identify where these differences occurred, a Bonferroni post hoc correction was applied, revealing that U16 consistently presented higher values than U13 across most variables (e.g., height: U16 vs. U13, p < 0.001; CMJ: U16 vs. U13, p < 0.001), with intermediate differences between adjacent groups (e.g., U14 vs. U15, p < 0.05 for CMJ).

In Table 2, the results of the Pearson correlation between the research variables were calculated separately for each age group (U13, U14, U15, U16) rather than for the entire sample because of significant intergroup differences. For each group, height was correlated with weight, sitting height, and leg length; weight was correlated with sitting height and leg length. Additionally, both height and weight were correlated with skeletal age, minutes of playing, football training, and CMJ but not with GH or IGF-1. Minute play was correlated with all variables except IGF-1 in all groups, and skeletal age followed a similar pattern. These correlations are detailed in Tables 3-6.

**Table 2:** Multiple comparisons between groups (Bonferroni post hoc correction)

| Variables | U13 vs U14 |  | U13 vs U15 |  | U13 vs U16 |  | U14 vs U15 |  | U14 vs U16 |  | U15 vs U16 |  |
| --- | --- | --- | --- | --- | --- | --- | --- | --- | --- | --- | --- | --- |
|  | p | ES | p | ES | p | ES | p | ES | p | ES | p | ES |
| Height (cm) | <0.001 | -1.44 | <0.001 | -1.79 | <0.001 | -2.43 | 1.000 | -0.35 | 0.003 | -0.96 | 0.175 | -0.61 |
| Weight (kg) | <0.001 | -1.23 | <0.001 | -1.59 | <0.001 | -1.83 | 0.345 | -0.44 | 0.062 | -0.62 | 1.000 | -0.16 |
| Sitting Height (cm) | <0.001 | -1.47 | <0.001 | -2.39 | <0.001 | -3.62 | <0.001 | -1.18 | <0.001 | -2.31 | 0.007 | -1.07 |
| Leg Length (cm) | <0.001 | -1.09 | 0.020 | -0.71 | 0.001 | -0.96 | 1.000 | 0.34 | 1.000 | -0.23 | 1.000 | -0.23 |
| Skeletal Age (yrs) | <0.001 | -2.07 | <0.001 | -1.89 | <0.001 | -4.06 | 1.000 | -0.35 | <0.001 | -2.07 | <0.001 | -1.72 |
| Football Training (mo) | 1.000 | -0.20 | 0.009 | -0.79 | 0.011 | -0.78 | 0.099 | -0.59 | 0.116 | -0.59 | 1.000 | 0.02 |
| $VO_{2max}$ (ml·kg<sup>-1</sup>·min<sup>-1</sup>) | <0.001 | -1.52 | <0.001 | -2.58 | <0.001 | -2.58 | 0.124 | -0.61 | 0.001 | -0.84 | 0.823 | -0.33 |
| CMJ (cm) | 0.139 | -0.65 | <0.001 | -2.10 | <0.001 | -2.65 | <0.001 | -1.43 | <0.001 | -1.92 | 1.000 | -0.27 |
| GH (ng/dl) | 1.000 | -0.31 | 1.000 | -0.29 | 0.170 | -0.59 | 1.000 | -0.10 | 1.000 | -0.22 | 1.000 | -0.34 |
| IGF1 (ng/dl) | 1.000 | -0.14 | 0.326 | -0.47 | 1.000 | 0.27 | 1.000 | -0.37 | 0.625 | 0.45 | 0.021 | 0.92 |
| Minutes of Playing (min) | <0.001 | -1.65 | <0.001 | -5.35 | <0.001 | -5.52 | <0.001 | -3.58 | <0.001 | -3.72 | 1.000 | -0.09 |

Abbreviations: $VO_{2max}$ : maximal oxygen consumption; CMJ: countermovement jump; GH: growth hormone; IGF1: insulin-like growth factor 1; Yrs: years; Mo: months.

**Table 3:** Pearson correlation between variables for the U13 Group (n=30)

| Variables | Height | Weight | Sit Height | Leg Length | Skel Age | Football Trn | VO2max | CMJ | GH | IGF1 | Min Playing |
| --- | --- | --- | --- | --- | --- | --- | --- | --- | --- | --- | --- |
| Height (cm) | 1 |  |  |  |  |  |  |  |  |  |  |
| Weight (kg) | .831** | 1 |  |  |  |  |  |  |  |  |  |
| Sitting Height (cm) | .875** | .793** | 1 |  |  |  |  |  |  |  |  |
| Leg Length (cm) | .905** | .695** | .585** | 1 |  |  |  |  |  |  |  |
| Skeletal Age (yrs) | .593** | .579** | .489** | .563** | 1 |  |  |  |  |  |  |
| Football Training (mo) | -.036 | .010 | .023 | -.080 | -.018 | 1 |  |  |  |  |  |
| $VO_{2max}$ (ml·kg<sup>-1</sup>·min<sup>-1</sup>) | .126 | -.113 | .053 | .165 | .056 | .067 | 1 | | | | |
| CMJ (cm) | .135 | .023 | .091 | .145 | .271 | .202 | .018 | 1 |  |  |  |
| GH (ng/dl) | .212 | .157 | .105 | .262 | .231 | -.200 | .225 | -.040 | 1 |  |  |
| IGF1 (ng/dl) | .406* | .238 | .487** | .251 | .344 | .177 | .118 | .344 | .106 | 1 |  |
| Minutes Playing (min) | .124 | -.134 | .050 | .164 | -.104 | -.073 | -.051 | .345 | -.194 | .137 | 1 |

**Table 4:** Pearson correlation between variables for the U14 group (n=31)

| Variables | Height | Weight | Sit Height | Leg Length | Skel Age | Football Trn | VO2max | CMJ | GH | IGF1 | Min Playing |
| --- | --- | --- | --- | --- | --- | --- | --- | --- | --- | --- | --- |
| Height (cm) | 1 |  |  |  |  |  |  |  |  |  |  |
| Weight (kg) | .824** | 1 |  |  |  |  |  |  |  |  |  |
| Sitting Height (cm) | .846** | .830** | 1 |  |  |  |  |  |  |  |  |
| Leg Length (cm) | .894** | .624** | .517** | 1 |  |  |  |  |  |  |  |
| Skeletal Age (yrs) | -.068 | .075 | .045 | -.146 | 1 |  |  |  |  |  |  |
| Football Training (mo) | -.290 | -.090 | -.078 | -.400* | .039 | 1 |  |  |  |  |  |
| VO<sub>2</sub>max (ml·kg<sup>-1</sup>·min<sup>-1</sup>) | -.416* | -.378* | -.280 | -.432* | .058 | .042 | 1 |  |  |  |  |
| CMJ (cm) | -.021 | .047 | .126 | -.140 | .308 | -.138 | .404* | 1 |  |  |  |
| GH (ng/dl) | -.321 | -.243 | -.398* | -.181 | -.099 | .032 | .093 | -.318 | 1 |  |  |
| IGF1 (ng/dl) | .295 | .322 | .329 | .198 | .033 | -.245 | .014 | .320 | -.006 | 1 |  |
| Minutes Playing (min) | -.236 | .036 | -.112 | -.285 | .348 | .216 | .106 | .102 | .266 | -.019 | 1 |

**Table 5:** Pearson correlation between variables for the U15 group (n=30)

| Variables | Height | Weight | Sit Height | Leg Length | Skel Age | Football Trn | VO2max | CMJ | GH | IGF1 | Min Playing |
| --- | --- | --- | --- | --- | --- | --- | --- | --- | --- | --- | --- |
| Height (cm) | 1 |  |  |  |  |  |  |  |  |  |  |
| Weight (kg) | .834** | 1 |  |  |  |  |  |  |  |  |  |
| Sitting Height (cm) | .865** | .776** | 1 |  |  |  |  |  |  |  |  |
| Leg Length (cm) | .941** | .748** | .645** | 1 |  |  |  |  |  |  |  |
| Skeletal Age (yrs) | .689** | .465** | .622** | .631** | 1 |  |  |  |  |  |  |
| Football Training (mo) | .015 | .093 | .065 | -.020 | -.003 | 1 |  |  |  |  |  |
| VO<sub>2</sub>max (ml·kg<sup>-1</sup>·min<sup>-1</sup>) | -.181 | -.176 | -.087 | -.217 | -.024 | .093 | 1 |  |  |  |  |
| CMJ (cm) | .311 | .182 | .379* | .219 | .383* | -.057 | .457* | 1 |  |  |  |
| GH (ng/dl) | -.095 | -.056 | -.055 | -.107 | .193 | -.036 | .481** | .301 | 1 |  |  |
| IGF1 (ng/dl) | .093 | .025 | -.007 | .147 | .020 | .130 | .239 | .305 | .167 | 1 |  |
| Minutes Playing (min) | .111 | .065 | .009 | .164 | .344 | .348 | .237 | .158 | .358 | .387* | 1 |

**Table 6:** Pearson correlation between variables for the U16 group (n=31)

| Variables | Height | Weight | Sit Height | Leg Length | Skel Age | Football Trn | VO2max | CMJ | GH | IGF1 | Min Playing |
| --- | --- | --- | --- | --- | --- | --- | --- | --- | --- | --- | --- |
| Height (cm) | 1 |  |  |  |  |  |  |  |  |  |  |
| Weight (kg) | .617** | 1 |  |  |  |  |  |  |  |  |  |
| Sitting Height (cm) | .721** | .264 | 1 |  |  |  |  |  |  |  |  |
| Leg Length (cm) | .921** | .669** | .394* | 1 |  |  |  |  |  |  |  |
| Skeletal Age (yrs) | .069 | .425* | .116 | .026 | 1 |  |  |  |  |  |  |
| Football Training (mo) | -.153 | -.146 | -.206 | -.087 | -.027 | 1 |  |  |  |  |  |
| VO<sub>2</sub>max (ml·kg<sup>-1</sup>·min<sup>-1</sup>) | -.449* | -.390* | -.438* | -.349 | -.298 | .069 | 1 |  |  |  |  |
| **CMJ (cm)** | -.321 | -.280 | -.156 | -.338 | -.049 | .054 | .400* | 1 |  |  |  |
| **GH (ng/dl)** | .150 | .073 | .172 | .102 | -.070 | .110 | .004 | .249 | 1 |  |  |
| **IGF1 (ng/dl)** | -.008 | -.128 | .044 | -.035 | -.285 | -.110 | .363* | .175 | .057 | 1 |  |
| **Minutes Playing (min)** | -.321 | -.358* | -.336 | -.237 | -.419* | -.024 | .570** | .399* | .285 | .368* | 1 |

Note: \*\* $p < 0.05$ , \* $p < 0.01$ . Correlations were calculated separately for each group (U13, $n=30$ ; U14, $n=31$ ; U15, $n=30$ ; U16, $n=31$ ) via Pearson's $r$ .

## Discussion

The present study compared hormone levels, physical fitness, skeletal age, and minutes of play in U13-16 elite football players and analyzed the relationships among all the variables. Research has shown that with the onset of puberty in players, their size and body structure change [29], and in this study, a similar change was observed. Significant differences in the anthropometric measurements were observed between the groups. The results revealed increases in height, weight, sitting height, and leg length as chronological age increased.

These between-group differences extend our earlier work on this population. Previous studies examined single age categories, under-15 or under-16 teams of 23 to 26 players, and focused on predicting individual playing time [25-28]. The present study instead compared four consecutive age groups (under-13 to under-16; n = 122) within one academy under a common protocol, allowing the maturational, hormonal, and fitness profile of each category to be positioned along a developmental continuum rather than examined in isolation. The progressive between-group increases in skeletal age, VO_2_max, CMJ, and IGF-1, together with the absence of a GH difference, describe how these characteristics diverge across the under-13 to under-16 transition, a pattern the single-age analyses could not reveal.

The results revealed that players in this academy were taller on average than their European counterparts, playing in Belgium [46], Portugal [47], and England [48]. However, they were also found to have similar characteristics to elite-level French clubs [49].

The physiological demands of football play are multifactorial [50]. The present study investigated measurements of physical fitness through the CMJ and VO_2_max (via YIRT1). Previously, both variables were shown to facilitate the prediction of progression in an academic setting [51]. Moreover, Le Gall et al. suggested that CMJ may help identify higher standards of football in international youth players in the U14 and U16 age groups [31], and Kelly et al. stated that this test should be included in all battery tests of football academies [52].

The increase in high-intensity running witnessed in the game over the last few seasons [53] highlights the importance of physical fitness, as assessed by CMJ and YIRT1 in this study. The CMJ is a reliable measure and relatively simple to perform. Despite the lack of diversity between the U11, U12, and U13 teams in Williams’ research [54], in this study, a significant difference was observed between all age groups, especially between U13 and U14, U13 and U15, U13 and U16, and U14 and U15. The results revealed significant differences in CMJ height with increasing age. The large difference when comparing those age categories suggests that maturation may have an impact on player development.

The VO_2_max analysis by YIRT1 showed a progressive and significant difference across each group, where older groups (e.g., U16) covered more distance in the test than younger ones did (e.g., U13). Previous research has shown that aerobic capacity is highly important to coaches at all academic levels [55]. The majority of football matches are dominated by high-intensity movements [55]. Not only the ability to achieve these speeds but also the ability to produce them repeatedly without hindering performance is crucial. Similar improvements in YIRT1 performance across chronological age groups were reported by Deprez et al. [56]. A possible reason for this distinct ability in older groups could be that older teams play 90 minutes per game, which could help develop aerobic capacity so that players can run longer before fatigue begins.

In addition, as players grow older, they develop more efficient movement patterns, which means that older players adapt better to changing directions than younger players do. Older players can also control their running speed, which allows them to improve their speed, which can account for the difference in YIRT1. For those reasons, it was proposed that growth and maturation data should be collected to provide effective training load and strength and conditioning training programs [52].

Another study has examined jump height in elite youth football players [54]. Specifically, the players jumped lower in CMJ tests than did the elite players within this academy. The players in this study were greater in terms of the height-to-weight ratio, which shows that they were able to achieve higher scores in CMJ.

Exercise is known to play an important role in regulating the GH-IGF-1 axis [57]. The GH response to exercise depends on the duration and intensity of exercise, the fitness of the subject, and environmental factors such as ambient temperature [58]. During adolescence, GH also regulates body composition and physiological responses to exercise [59]. In this study, GH levels did not significantly differ across groups. Given the high inter-individual variability of GH and the resulting limited statistical power, this should be interpreted as the absence of a detectable between-group difference in the present sample rather than as evidence that age or football training does not affect GH.

Serum IGF-1 levels increase steadily at the beginning of puberty, peak in late puberty, and then decrease rapidly [60]. Contradictory results have been reported regarding the effect of exercise on serum IGF-1 [14]. As with our GH results, our findings of increasing IGF-I levels with age are not consistent with those reported by Mejri et al. [17]. In young people, high-intensity exercise is associated with increased IGF activity in favor of anabolic status [14]. The results of the present study revealed that with increasing age in elite players, the levels of these hormones also increase. There seems to be a need for more research on these hormones and different age groups before and after puberty to better determine whether the levels of these hormones increase with increasing exercise and age or later decline from puberty.

Using anthropometric-based predictive equations for skeletal age and tests such as the medicine ball throw and vertical jump for upper and lower limb strength, Gantois et al. examined the association between explosive strength and skeletal age in 239 young volleyball players (both sexes, ages 10–13). Regardless of sex, they discovered a positive correlation between skeletal age and explosive strength, with early-maturing players but late-maturing players having lower [32]. However, these findings contrast with the present results since skeletal age did not correlate with any of the physical tests at U13. The differences in sample size and sports (volleyball versus football) may explain the contrasting results. Moreover, the current study revealed that skeletal age was positively correlated with CMJ performance in the U15 group (r = 0.383, p < 0.05) and that there were significant differences between age groups (p < 0.001) among elite male football players (U13–U16). The highest CMJ values were found in U16 players (40.87 ± 3.87 cm), in contrast to U13 players (29.67 ± 3.48 cm). However, unlike Gantois et al., our work assessed skeletal age via the Fels method, which might be more accurate than anthropometric projections and could account for the larger group differences observed. These results demonstrate how biological maturation affects young athletes’ explosive power, but the necessity for sport-specific research is highlighted by our focus on male football players and various approaches to assessment.

The correlation analysis revealed various relationships across groups (Tables 3-6). In U13, height was strongly correlated with sitting height (r = 0.875, p < 0.01) and leg length (r = 0.905, p < 0.01) but only moderately correlated with IGF-1 (r = 0.406, p < 0.05). In U14, height had a moderate negative correlation with VO_2_max (r = -0.416, p < 0.05), suggesting that taller players may have lower aerobic capacity at this age. U15 and U16 showed fewer consistent correlations, with U16 displaying a strong negative correlation between minutes of playing and skeletal age (r = -0.419, p < 0.05). The lack of consistent correlations with GH and IGF-1 across all groups indicates that hormonal responses may be influenced by factors beyond anthropometric and fitness variables, such as individual maturation timing [26]. However, these findings contrast with those of a previous study that reported that players with greater playing time presented higher values of both CMJ and IGF-1 [25]. Moreover, the present study did not find any associations between VO_2max_ and GH, GH or CMJ, or between playing time and CMJ, which contrasts with the findings of another study [28]. In addition, other associations between VO_2_max and CMJ or playing time were not evident, which contrasts with the results of previous studies [25]. Even so, it is relevant to mention the differences in the sample size and the context of the within-team analysis that may explain the results.

### Limitations

This study has several limitations. First, the cross-sectional design limits the ability to infer causality between variables such as training duration and hormonal changes. Second, the sample was drawn from a single academy in Iran, which may not represent other populations or training environments. Third, GH and IGF-1 measurements were taken at a single time point, potentially missing diurnal or exercise-induced fluctuations. Finally, the study did not account for dietary factors or sleep patterns, which could influence hormonal and physical outcomes.

### Practical Implications

These findings suggest that coaches should prioritize biological maturity over chronological age when designing training programs for youth football players. Tailoring training loads to match the skeletal age could optimize physical development and reduce the degree of injury risk. Additionally, the significant differences in CMJ and VO_2_max highlight the importance of incorporating explosive power and aerobic capacity tests in talent identification processes.

### Suggestions for Future Research

Future studies should adopt longitudinal designs to track changes in hormonal, anthropometric, and fitness variables over time. Investigating the effects of diet, sleep, and training intensity on GH and IGF-1 responses could provide deeper insights. Additionally, including female players and diverse populations would increase generalizability. Finally, exploring the impact of early versus late maturation on long-term athletic success could inform talent development strategies.

## Conclusion

In conclusion, with age, the anthropometric characteristics and hormonal characteristics of elite players change. In terms of performance prediction, owing to the available data, this study can provide some clear insight. It is recommended that clubs compare players in terms of maturity rather than relative or chronological age. As a result, there is the potential for the Academy to be able to more effectively protect younger players and allow them to grow according to their normal growth before releasing them and reaching their physical potential.

This requires a change in the way coaches think and scout about players and a focus on their technical ability rather than their physical competence.

## Declaration of generative AI and AI-assisted technologies in the writing process

During the preparation of this work, the author used Grok AI to improve language and readability. After using this tool, the author reviewed and edited the content as needed and takes full responsibility for the content of the publication.

